# The *Chlamydomonas reinhardtii* CLiP2 mutant collection expands genome coverage with high-confidence disrupting alleles

**DOI:** 10.1101/2024.12.16.626622

**Authors:** Alice Lunardon, Weronika Patena, Cole Pacini, Michelle Warren-Williams, Yuliya Zubak, Matthew Laudon, Carolyn Silflow, Paul Lefebvre, Martin Jonikas

**Affiliations:** Princeton University, NJ, USA; University of Minnesota, MN, USA; Howard Hughes Medical Institute

## Abstract

*Chlamydomonas reinhardtii* (Chlamydomonas hereafter) is a powerful model organism for studies of photosynthesis, ciliary motility, and other cellular processes [1–4]. The CLiP library of mapped nuclear random insertion mutants [5,6] has accelerated progress for hundreds of laboratories in these fields by providing mutants in genes of interest. However, its value was limited by its modest coverage of the genome with high-confidence disruption alleles (46% of nuclear protein-coding genes with 1+ high-confidence allele in exons/introns; 12% of genes with 3+ alleles in exons/introns). Here we introduce the CLiP2 (Chlamydomonas Library Project 2) library, which greatly expands the number of available mapped high-confidence insertional mutants. The CLiP2 library includes 71,700 strains, covering 79% of nuclear protein-coding genes with 1+ high-confidence allele in exons/introns and 49% of genes with 3+ alleles in exons/introns. The mutants are available to the community via the Chlamydomonas Resource Center.

## Introduction

The CLiP2 mutants have many similarities to the original CLiP mutants, with several noteworthy differences. Specifically, the mutants were generated in a different background strain, of a mating type opposite to the original collection, with a different cassette that confers a different antibiotic resistance. Below, we provide more information on these differences, data on genome coverage, and guidance for use of the mutants.

### Background strain

The background strain for CLiP2 mutants is CC-5415, a mating type plus (mt+) Chlamydomonas strain carrying the *nit1 agg1* mutations, also known as Witman g1 [7]. CC-5415 was chosen for the CLiP2 library due to the following characteristics: 1) it exhibits robust negative phototaxis and swims well, important for studies on phototaxis and motility; 2) CC-5415 (mt+) has the opposite mating type and mates efficiently with the parental strain of the original CLiP mutants (CMJ030 = CC-4533), facilitating the generation of double mutants via genetic crosses; 3) CC-5415 has a cell wall, which is beneficial for certain studies; and 4) we observed that it replicates well robotically, which is essential for propagation of a large mutant collection.

### DNA cassette

We generated CLiP2 mutants by random genomic insertion of the CIB2 DNA cassette, which confers hygromycin resistance. This is in contrast to the CIB1 cassette used to generate CLiP1 mutants, which confers paromomycin resistance. We chose a different antibiotic marker to facilitate the selection of double mutants when crossing CLiP and CLiP2 strains.

Like CIB1, each CIB2 cassette normally contains two unique 22-nucleotide barcodes, one at the 5’ end and one at the 3’ end of the cassette, allowing for unique identification of each insertion.

Expression of the AphVII gene, conferring resistance to hygromycin, is driven by two promoters, the heat shock protein 70A (hsp70A) and the beta 2-tubulin (tubB2), and terminated by two terminators, Photosystem I subunit (psad) and ribosomal protein 12 (rpl12), oriented in opposite directions. The intent behind these two terminators is that when the cassette inserts into a gene, one of the terminators will be in the same orientation as the gene and will terminate the gene’s transcript, resulting in a truncated protein and degradation of the mRNA by nonsense-mediated decay. The full sequence of the CIB2 cassette (with the unique barcode sequences indicated with Ns) is as follows:

ATGACTCCACGAGTGTCGCTGACCGTGGATGCTCAGTAGTCACACGAGCCCTCGTCAGAAACACGTCTCCNNNNNNNNNNNNNNNNNNNNNNGGCAAGCTAGAGAACCATCCATCAGGCAGCTCGCTGAGGCTTGACATGATTGGTGCGTATGTTTGTATGAAGCTACAGGACTGATTTGGCGGGCTATGAGGGCGGGGGAAGCTCTGGAAGGGCCGCGATGGGGCGCGCGGCGTCCAGAAGGCGCCATACGGCCCGCTGGCGGCACCCATCCGGTATAAAAGCCCGCGACCCCGAACGGTGACCTCCACTTTCAGCGACAAACGAGCACTTATACATACGCGACTATTCTGCCGCTATACATAACCACTCAGCTGGCTTAAGATCCCATCAGGCTTGCATGGCAGCCAAACCAGGATGATGACTTGCAACCCTTATCCGGAATCAAGCTTCTTTCTTGCGCTATGACACTTCCAGCAAAAGGTAGGGCGGGCTGCGAGACGGCTTCCCGGCGCTGCATGCAACACCGATGATGCTTCGACCCCCCGAAGCTCCTTCGGGGCTGCATGGGCGCTCCGATGCCGCTCCAGGGCGAGCGCTGTTTAAATAGCCAGGCCCCCGATTGCAAAGACATTATAGCGAGCTACCAAAGCCATATTCAAACACCTAGATCACTACCACTTCTACACAGGCCACTCGAGCTTGTGATCGCACTCCGCTAAGGGGGCGCCTCTTCCTCTTCGTTTCAGTCACAACCCGCAAACATGACACAAGAATCCCTGTTACTTCTCGACCGTATTGATTCGGATGATTCCTACGCGAGCCTGCGGAACGACCAGGAATTCTGGGAGGTGAGTCGACGAGCAAGCCCGGCGGATCAGGCAGCGTGCTTGCAGATTTGACTTGCAACGCCCGCATTGTGTCGACGAAGGCTTTTGGCTCCTCTGTCGCTGTCTCAAGCAGCATCTAACCCTGCGTCGCCGTTTCCATTTGCAGCCGCTGGCCCGCCGAGCCCTGGAGGAGCTCGGGCTGCCGGTGCCGCCGGTGCTGCGGGTGCCCGGCGAGAGCACCAACCCCGTACTGGTCGGCGAGCCCGGCCCGGTGATCAAGCTGTTCGGCGAGCACTGGTGCGGTCCGGAGAGCCTCGCGTCGGAGTCGGAGGCGTACGCGGTCCTGGCGGACGCCCCGGTGCCGGTGCCCCGCCTCCTCGGCCGCGGCGAGCTGCGGCCCGGCACCGGAGCCTGGCCGTGGCCCTACCTGGTGATGAGCCGGATGACCGGCACCACCTGGCGGTCCGCGATGGACGGCACGACCGACCGGAACGCGCTGCTCGCCCTGGCCCGCGAACTCGGCCGGGTGCTCGGCCGGCTGCACAGGGTGCCGCTGACCGGGAACACCGTGCTCACCCCCCATTCCGAGGTCTTCCCGGAACTGCTGCGGGAACGCCGCGCGGCGACCGTCGAGGACCACCGCGGGTGGGGCTACCTCTCGCCCCGGCTGCTGGACCGCCTGGAGGACTGGCTGCCGGACGTGGACACGCTGCTGGCCGGCCGCGAACCCCGGTTCGTCCACGGCGACCTGCACGGGACCAACATCTTCGTGGACCTGGCCGCGACCGAGGTCACCGGGATCGTCGACTTCACCGACGTCTATGCGGGAGACTCCCGCTACAGCCTGGTGCAACTGCATCTCAACGCCTTCCGGGGCGACCGCGAGATCCTGGCCGCGCTGCTCGACGGGGCGCAGTGGAAGCGGACCGAGGACTTCGCCCGCGAACTGCTCGCCTTCACCTTCCTGCACGACTTCGAGGTGTTCGAGGAGACCCCGCTGGATCTCTCCGGCTTCACCGATCCGGAGGAACTGGCGCAGTTCCTCTGGGGGCCGCCGGACACCGCCCCCGGCGCCTGATCTAGATGGCAGCAGCTGGACCGCCTGTACAATGGAGAAGAGCTTTACTTGCCGGGATGGCCGATTTCGCTGATTGATACGGGATCGCAGCTCGGAGGCTTTCGCGCTAGGGGCTAGGCGAAGGGCAGTGGTGACCAGGGTCGGTGTGGGGTCGGCCCACGGTCAATTAGCCACAGGAGGATCAGGGGGAGGTAGGCACGTCGACTTGGTTTGCGACCCCGCAGTTTTGGCGGACGTGCTGTTGTAGATGTTAGCGTGTGCGTGAGCCAGTGGCCAACGTGCCACACCCATTGAGAAGACCAACCAACTTACTGGCAATATCTGCCAATGCCATACTGCATGTAATGGCCAGGCCATGTGAGAGTTTGCCGTGCCTGCGCGCGCCCCGGGGGCGGAATCTGCTGCCGGCTGCCCCCGCCCCCGCTGCTGAGGCTGAATCTACTCCGCTCTCCACCCCAGTCCCAGGAAGAGAGGCGCATTTACTTCGCACAGACGTTACAGCACACCCTTGATCATCATCAGCTGCTCTTCCCTGCCGCTGCAACACGCCCCGCGCTANNNNNNNNNNNNNNNNNNNNNNACTGACGTCGAGCCTTCTGGcAGACTGCTACGATGACCGACACTGCGGACCTCGAACTGGATTCAG

### Transformation protocol

The following protocol was used to generate the CLiP2 mutants: the Chlamydomonas CC-5415 strain was grown in TAP medium [8] in a 500mL flask under 100 μmol photons m^-2^ s^-1^ light to a density of 0.5-2 × 10^6 cells/mL. Cells were collected by centrifugation at 1000 x g for 5 min in batches of 50 mL. Pellets were washed twice with 2.5 mL GeneArt® MAX Efficiency® Transformation Reagent for Algae (Thermo Fisher Scientific), and then resuspended in GeneArt® MAX Efficiency® Transformation Reagent for Algae at 2-3 x 10^8 cells/mL. 50 µL cell suspensions were then aliquoted in pre-chilled 1.5 mL tubes. For each tube, 5ng of CIB2 DNA at 5 ng/µL was added to the cell suspension and mixed by tapping the tube. The mixture was then transferred to a pre-chilled 2mm electroporation cuvette (12358-346, Bulldog-Bio). Electroporation was immediately performed with a NEPA21 electroporator (Nepa Gene) with the following settings: poring pulse: voltage: 300V; pulse length: 8 msec; pulse interval: 50 msec; number of pulses: 2; decay rate: 40%; polarity: +; transfer pulse: voltage: 20V; pulse length: 50 msec; pulse interval: 50 msec; number of pulses: 5; decay rate: 40%; polarity: +/-. For ∼20% of the transformations reactions, the poring pulse decay rate was set to 10%, instead of 40%, and the transfer pulse number of pulses was set to 1, instead of 5. After electroporation, the cells in the cuvette were incubated at room temperature for 15 min and then transferred to a 15mL tube containing 8 mL TAP supplemented with 40 mM sucrose and shaken gently in the dark for 6 h. After incubation, cells were collected by centrifugation at 1000 x g for 5 min and resuspended in 1 mL TAP. Cells were then plated (250 µL per plate) on rectangular PlusPlates™ (Singer Instruments) containing ∼70mL TAP agar [8] with 20 µg/mL hygromycin and incubated in low light (<5 μmol photons m^-2^ s^-1^ light) for approximately two weeks before colony picking. Colonies were then robotically picked and arrayed at 384 colonies per plate as previously described [6].

### Mutant propagation

Mutants were propagated as arrays of 384 strains per plate on rectangular PlusPlates™ (Singer Instruments) containing agar. During the initial ∼20 replications while the mutants were being mapped at Princeton University, the agar used was TAP agar [8] with 20 µg/mL hygromycin. For subsequent replications after the collection was transferred to the Chlamydomonas Center, the agar used was (S)TAP agar [9] with 20 µg/mL hygromycin.

The plates were propagated with a ROTOR (Singer Instruments) with the following settings. Source pinning: 1) Pressure: 30%; 2) Speed: 5 mm/s. Source dry mix: 1) Clearance: 0.5mm; 2) Diameter: 1.2mm. Target pinning: 1) Pressure: 30%; 2) Speed: 5 mm/s. Target dry mix: 1) Clearance: 0.5mm; 2) Diameter: 1mm. For the first few replications, mutants were replicated with a Speed of 19 mm/s, but this was observed to occasionally lead to splattering of the colonies, so to reduce the incidence of cross-contamination, the speed was reduced to 5 mm/s. Mutants have been propagated in the same light conditions as used for selection.

### Insertion mapping

The insertion sites in the CLiP2 library were mapped using the pipeline used for the original CLiP library [6], with some improvements described below. This pipeline identified the barcodes in each mutant colony and determined the genomic DNA sequences flanking each barcoded insertion. We implemented two noteworthy improvements to the pipeline. One major improvement was that much longer (up to 6 kb instead of up to 1 kb) Leap-Seq products were generated and sequenced by Pacific Biosciences long-read sequencing instead of Illumina sequencing, allowing us to read past most “junk” DNA fragments, which are major source of incorrect mappings. A second major improvement was that we read not only outward from each barcode but also inward across the cassette and into the flanking genomic DNA on the opposite side of the cassette, allowing further confirmation of the genomic insertion site in mutants with a truncated cassette, and further improving our mapping accuracy.

Insertions were considered to be mapped with high confidence if we mapped both sides of the insertion to the same locus, or if we only mapped one side but our analysis flagged them as high-quality, meaning data reading at least 900bp into the genome, and 2+ reads confirming the position, constituting over 50% of all reads for the given insertion junction.

### Mutant cherry-picking

Our insertion mapping yielded 197,703 mutants with mapped insertions, from which we cherry-picked mutants to reduce the collection size to make it manageable for long-term propagation and storage. We selected for cherry-picking all mutants with insertions in genes, up to a maximum of 5 mutants with high mapping confidence per gene, or up to 5 additional mutants if the top 5 mutants were not mapped with high confidence, to increase the chance of including 5 correctly-mapped mutants per gene. When a gene was represented by more than 5 mutants, we prioritized mutants by a combination of mapping confidence and likelihood of disrupting the gene (whether the insertion is in an exon/intron or in the 5’ or 3’ UTR, and how central the insertion position is in the gene sequence). We cherry-picked mutants using a PIXL robot (Singer Instruments) to 190 plates in 384-colony format, for a final collection of 71,700 strains. Mutant insertion sites in the final cherry-picked collection are shown in Table S1.

### Genome coverage

We characterized the genome coverage according to four categories of insertions: all insertions, intron/exon insertions, insertions with high mapping confidence, and high-confidence intron/exon insertions (Figure 1). When selecting mutants, we recommend prioritizing insertions represented in the last category (high confidence exon/intron insertions) for functional analyses because they are the most likely to disrupt gene expression and yield observable phenotypes. Altogether, the final collection of 71,700 strains covers 79% of nuclear protein-coding genes with 1+ high-confidence allele in exons/introns and 49% of genes with 3+ alleles in exons/introns.

**Figure 1.**
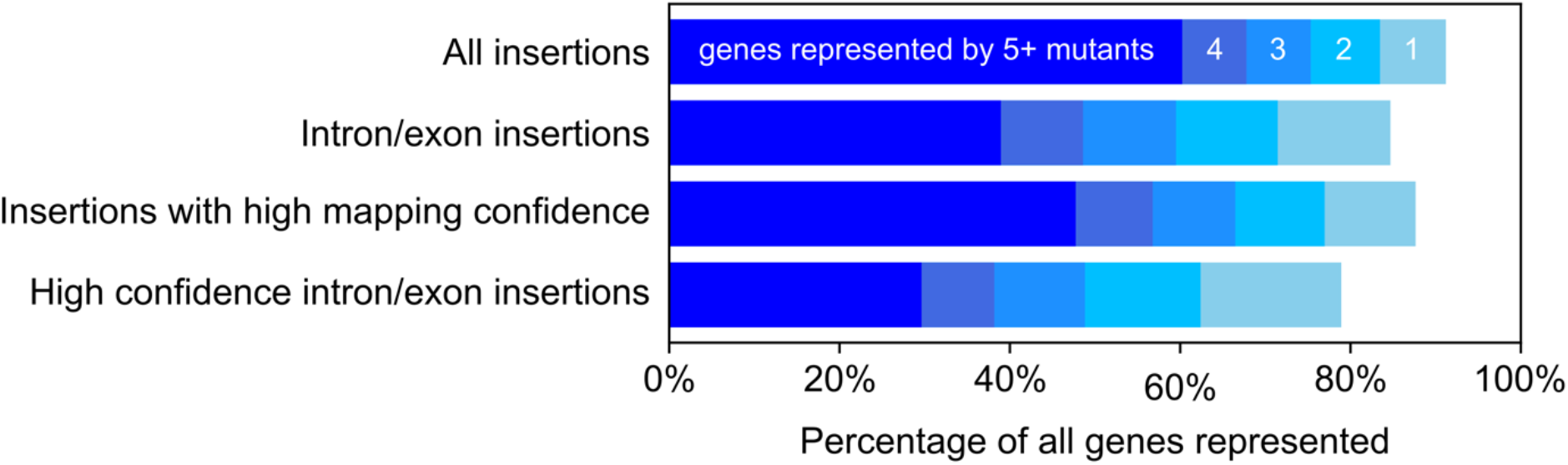
The CLiP2 library covers most of the genome with multiple high-confidence alleles. Genome coverage is shown for four different categories of insertions. The top bar shows the total genome coverage, encompassing all insertions in any gene feature (exons, introns, untranslated regions [UTRs]) and irrespective of mapping confidence levels (both low and high). The second bar focuses on insertions specifically in exons or introns, excluding UTR insertions, which are less likely to disrupt gene function and produce phenotypes [6]. The third bar restricts coverage to insertions that were mapped with high confidence, ensuring greater accuracy in identifying the mutation site. The bottom bar combines both restrictions, showing insertions in exons or introns mapped with high confidence. Within the bar representing each category, the data are visualized using different shades of blue, corresponding to the number of distinct mutant alleles available for each gene. The lightest blue section on the right of each bar indicates the percentage of genes with only one available mutant allele, transitioning through progressively darker shades to the darkest blue section on the left, which represents genes with five or more independent mutant alleles.

### Mutants can be searched and ordered online

To search for mutants with insertions in genes of interest, please visit the CLiP website (https://www.chlamylibrary.org/). The CLiP website also includes links to order the mutants from the Chlamydomonas Resource Center (https://www.chlamycollection.org/).

### Need for streaking to single colonies and validation of insertion sites by PCR

The reported insertions have a chance of being incorrectly mapped. Furthermore, due to the limitations of the robotic methods used for propagation, some mutants may be contaminated with other mutants. Therefore, as soon as you receive your mutant and before investing further effort into characterizing it, we urge you to streak the mutant to single colonies and confirm that your gene of interest is disrupted by using the PCR protocol available on the CLiP website (https://www.chlamylibrary.org/files/Instructions%20on%20PCRs%20to%20check%20the%20insertion%20site.pdf).

### Need for validation of genotype-phenotype relationship

Please also be aware that, as with previous collections, there may be additional unmapped insertions and/or mutations in the genome of any given mutant. One therefore cannot immediately conclude that a mutant’s phenotype is due to that mutant’s mapped insertion. To establish that the disruption of a particular gene causes a particular phenotype, we recommend attempting to genetically rescue the mutant’s phenotype by transforming the mutant with a wild-type copy of the gene via PCR amplification from the wild-type genome. Alternatively, backcrossing the mutant with the wild type and checking for co-segregation of the phenotype with hygromycin resistance can also verify the association. However, this method cannot exclude the possibility that the phenotype is actually caused by a mutation in a different gene near the cassette insertion.

## Discussion

We hope this resource will help accelerate progress in fields for which Chlamydomonas serves as a model system. We are grateful to the community for the support of this work.

## Acknowledgments

This work was supported by National Science Foundation grant MCB-1914989 and by the Howard Hughes Medical Institute. The Chlamydomonas Resource Center is funded by NSF award DBI 2247108. We would like to thank Matthew LaVoie and Savannah Briggs for help with maintaining and freezing the library at the CRC.

---

**Table S1. Insertion sites of the mutants in CLiP2**. Each line of the file corresponds to a single junction between the insertion cassette and the genome. Most insertions have two mapped junctions, some just have one. Multiple junctions in the same mutant can be two sides of the same insertion (most commonly when they’re mapped to the same locus), or multiple independent insertions in one mutant. The columns are as follows: mutant ID, side of cassette most proximal to the junction (5’ or 3’, with a note if we think the cassette may be truncated on this end, which can happen if the data are derived from the read inward across the cassette from the opposite side), chromosome, chromosome position, CreID of disrupted gene (if the position is part of multiple overlapping genes, they are separated by “&” in this field and all further fields related to genes), feature of disrupted gene, percentile position of disruption in the gene CDS, transcript names of the disrupted gene, the internal barcode sequence, the mapping confidence (very approximate and quite conservative), the suggested check PCR primers, and the Phytozome description of the disrupted gene.

## References

1. Lu H, Li Z, Li M, Duanmu D. Photosynthesis in Chlamydomonas reinhardtii: What We Have Learned So Far? In: Wang Q, ed. Microbial Photosynthesis. Springer Singapore; 2020:121–136. doi:10.1007/978-981-15-3110-1_6

2. Dupuis S, sMerchant SS. Chlamydomonas reinhardtii: a model for photosynthesis and so much more. Nat Methods. 2023;20(10):1441–1442. doi:10.1038/s41592-023-02023-6

3. Silflow CD, Lefebvre PA. Assembly and motility of eukaryotic cilia and flagella. Lessons from Chlamydomonas reinhardtii. Plant Physiol. 2001;127(4):1500–1507.

4. Marshall WF. Chlamydomonas as a model system to study cilia and flagella using genetics, biochemistry, and microscopy. Front Cell Dev Biol. 2024;12:1412641. doi:10.3389/fcell.2024.1412641

5. Li X, Zhang R, Patena W, et al. An Indexed, Mapped Mutant Library Enables Reverse Genetics Studies of Biological Processes in Chlamydomonas reinhardtii. Plant Cell. 2016;28(2):367–387. doi:10.1105/tpc.15.00465

6. Li X, Patena W, Fauser F, et al. A genome-wide algal mutant library and functional screen identifies genes required for eukaryotic photosynthesis. Nat Genet. 2019;51(4):627–635. doi:10.1038/s41588-019-0370-6

7. Pazour GJ, Sineshchekov OA, Witman GB. Mutational analysis of the phototransduction pathway of Chlamydomonas reinhardtii. J Cell Biol. 1995;131(2):427–440. doi:10.1083/jcb.131.2.427

8. Kropat J, Hong-Hermesdorf A, Casero D, et al. A revised mineral nutrient supplement increases biomass and growth rate in Chlamydomonas reinhardtii. Plant J. 2011;66(5):770–780. doi:10.1111/j.1365-313X.2011.04537.x

9. (S)TAP. https://www.chlamycollection.org/content/uploads/2021/03/STAP-Growth-Medium.pdf

